# Recurrent expansion and rapid evolution of the Drosophilid RNAi pathway in testis

**DOI:** 10.1101/2025.11.28.691189

**Authors:** Kimberly Elicker, Meysoon Quraishi, Garima Setia, Xin Yu Zhu Jiang, Lijuan Kan, Alex S. Flynt, Jiayu Wen, Lamia Wahba, Eric C. Lai

**Author notes:** These authors contributed equally (Kimberly Elicker, Meysoon Quraishi, Garima Setia).

## Abstract

Multiple classes of selfish genetic elements, including transposable elements, meiotic drivers, and viruses, are suppressed by small interfering RNAs (siRNAs) that guide RNA interference (RNAi). These are interlocked within continually evolving genetic conflicts that are characterized by extremely rapid dynamics of change in both sequence and copy number. Indeed, it was shown that across *Drosophila*, the core RNAi machinery evolves under positive selection, and that the RNAi effector *AGO2* has additional copies in some species. This contrasts with the core microRNA (miRNA) machinery in *Drosophila*, which evolves under negative selection and maintains one-to-one orthologs not only amongst flies, but even to mammals. Here, we analyze >300 long read genome assemblies of *Drosophila* species to generalize these attributes. Not only do we find recurrent expansion of *AGO2* across ∼100 species, including lineages with ongoing amplification of AGO2, we also find dozens of species with extra copies of the other core RNAi factors, *r2d2* and *dicer-2*. In many cases, these additional RNAi factor copies co-exist, or are nested, within a lineage bearing an ancestral expansion of *AGO2*. Transcriptome data provide evidence that core RNAi factors, including certain lineage-specific copies, are biased for testis expression. Finally, we use small RNA data to annotate hairpin RNAs (hpRNAs) in *D. pseudoobscura*, one of the species with prominent amplification of RNAi factors. We find rampant *de novo* hpRNA loci in this species, whose siRNAs are predominantly expressed in testis. All together, these findings highlight evolutionary plasticity of the fly RNAi pathway and affirm that it is preferentially deployed in the germline of the heterogametic sex. When considered alongside abundant genetic data for recurrent sex ratio meiotic drive against the Y chromosome, these genomic data strongly imply that a fundamental role of endogenous RNAi is to control sex chromosome conflicts.

## Introduction

Genomes are under constant attack, and require a multitude of robust defense mechanisms to prevent hijacking by selfish genetic elements. The most well-studied of these include transposable elements (TEs) and viruses. These collectively utilize vertical and horizontal modes of transmission to spread across populations, which highlight some of the challenges for their suppression, as well as strategies by which host systems might try to surveil and act specifically on these genomic intruders. Such defenses act at chromatin, RNA and protein levels, and sometimes work at multiple levels, highlighting mechanistic complexity as well as the strong biological imperative to innovate new defenses against relentless assaults.

Argonaute-based small RNA systems are particularly germane to suppression of selfish genetic elements. In many invertebrate species, three small RNA systems exist. (1) The microRNAs (miRNAs) are ∼22 nucleotide (nt) that are generally produced by RNase III enzymes (nuclear Drosha and cytoplasmic Dicer) and guide Ago-class effectors to endogenous target genes, which primarily bear short complementarity to positions 2-8 of the miRNA (the “seed” region) [1]. (2) Small-interfering RNAs (siRNAs) are ∼21 nt regulatory RNAs produced by a Dicer-class RNase III enzyme and loaded into specialized Ago proteins that harbor endoribonucleolytic cleavage activity (“Slicer” enzymes) [2]. siRNAs direct the cleavage of highly complementary targets, in a process generally referred to as RNA interference (RNAi). (3) Piwi-associated RNAs (piRNAs), which are ∼24-32 nt RNAs that populate PIWI-clade Argonautes that are frequently germline-restricted [3]. Of these pathways, siRNAs and piRNAs are critical to defense of selfish genetic elements. One of the most overt roles of siRNAs and RNAi is to combat viral infection, which can easily spread across individuals and sometimes between species. This role is shared by *Drosophila*, *C. elegans*, and plants, and may potentially have specialized utility in mammals (which lack a dedicated RNAi pathway, but do harbor the Slicer enzyme Ago2). The most well-understood role of piRNAs is to silence TEs in germlines, thereby suppressing their vertical amplification and limiting DNA damage in meiotic settings. This role is well-conserved between *Drosophila* and vertebrates, while a divergent piRNA pathway in *C. elegans* also safeguards the germline.

Most miRNAs that accumulate to substantial levels are substantially conserved, and are also associated with large cohorts of conserved seed matches across the transcriptome, particularly within 3’ UTRs [4]. On the other hand, the catalogs of abundant siRNAs and piRNAs from well-studied organisms are typically divergent within their closest relatives [5, 6], and can even be substantially polymorphic across individuals of a given population [7, 8]. In some cases, the small RNA sequences change, but they derive from syntenic transcription units. In other cases, there is *de novo* emergence or disappearance of small RNA progenitor loci themselves. The rapid turnover of well-expressed siRNAs and piRNAs is mirrored by their biological usage in suppression of viruses and TEs, which both exhibit extremely rapid evolutionary dynamics.

However, there is a third general class of selfish genetic element. Unlike TEs and viruses, which are nominally “non-self” (even if TEs are embedded in the genome), so-called meiotic drivers hijack Mendelian inheritance to enable their unfair transmission into productive gametes and/or progeny [9, 10]. This class of selfish element is particularly insidious, since they lack some of the characteristic features of TEs and viruses that are used to launch specific defenses whilst sparing host gene expression. In this case, meiotic drivers are “just genes”, and consequently largely escape the notice of humans studying the relevant genomes. However, they can have powerful effects on gametogenesis, by biasing their inheritance into >95% of progeny in some cases. While this can apply to autosomes [11, 12], there seems to be biased recurrence of meiotic drivers to live on a sex chromosome, and to disadvantage the heterogametic sex [13]. Thus, in XY systems, it is common for meiotic drivers to reside on the X chromosome and have activity specifically during spermatogenesis to disable Y-bearing sperm.

The molecular identities of such “sex ratio meiotic drivers” are mostly unknown, and even for the few systems cloned, their mechanisms of action are relatively opaque. Nevertheless, molecular genetic studies in Drosophila species highlight several related principles. First, in two species studied thus far (*D. melanogaster* [*Dmel*] and *D. simulans* [*Dsim*]), knockouts of core RNAi factors highlight preferential requirements in the male germline for either optimal spermatogenesis (*Dmel*) [14], or indeed outright male fertility (*Dsim*) [15]. Second, the testis-related phenotypes of RNAi mutants are substantially attributable to the loss of testis-restricted siRNAs from long inverted repeats known as hairpin RNAs (hpRNAs), which harbor specific, extensively complementary targets. In Dsim, some of these hpRNAs (e.g. Nmy and Tmy) directly repress professional X-linked sex ratio meiotic drivers of the Dox superfamily [15–19]. Additional de novo hpRNAs in Dsim (i.e., not found in Dmel or in other outgroup fly species) also target X-linked genes, which are candidates for novel sex ratio meiotic drivers [5]. Third, RNAi-related genes such as the effector *AGO2* evolve rapidly [20], akin to more conventional immunity genes [21], and presumably reflects their role in containing genomic conflicts. Notably, there are also extra copies of *AGO2* genes in several Dipterans [22], including within the obscura clade of Drosophilids [23–25], and many of these are testis-restricted. Altogether, these observations shine a light on fast-evolving hpRNA-siRNA loci and expansion of Ago2 paralogs in testis, which conceivably may reflect shared genetic utilization.

In this study, we take advantage of recent availability of >300 nanopore genomes of *Drosophila* species [26, 27], which enables studies of the dynamics of small RNA factor evolution on a much larger scale than previously possible. While core miRNA factors are generally stable, we find numerous cases where RNAi machinery has been amplified, including *AGO2*, *dicer-2* and *r2d2* genes. We analyze existing and newly made transcriptome data to provide general evidence for male and/or testis-biased expression of de novo RNAi gene copies. The most rampant amplification of RNAi factors occurs in the obscura clade. Concurrent with this, we uncover scores of *de novo*, testis-biased, hpRNAs in *D. pseudoobscura*. Altogether, this work provides strong evolutionary support to the notion that recurrent bouts of meiotic drive activity in the male germline are countered by amplification of RNAi activity and emergence of hpRNA loci.

## Results

### Annotation of core miRNA factors across 300+ Drosophilid genomes

The recent efforts of Kim and Petrov released hundreds of long read genome assemblies of *Drosophilidae* family species, providing a new foundation for assessment of molecular evolution across this genus [26, 27]. In addition to the bulk of species in the *Drosophila* genus, this dataset includes a selection of more distant related fruitflies in the *Zaprionus*, *Lordiphosa*, *Chymomyza* genera, amongst others. While a powerful dataset, a prominent limitation is that they are not currently annotated, and transcriptome evidence is lacking for most species. While this work was in progress, Obbard and colleagues made draft annotations *ab initio* via a multistep pipeline [28]. This framework works well for conserved genes, but rapidly evolving and/or *de novo* genes present substantial challenges to automatic pipelines. Therefore, we elected to conduct systematic tblastn searches across the genomes, followed by extensive manual inspection and curation of the hits (**Figure 1**).

**Figure 1.**
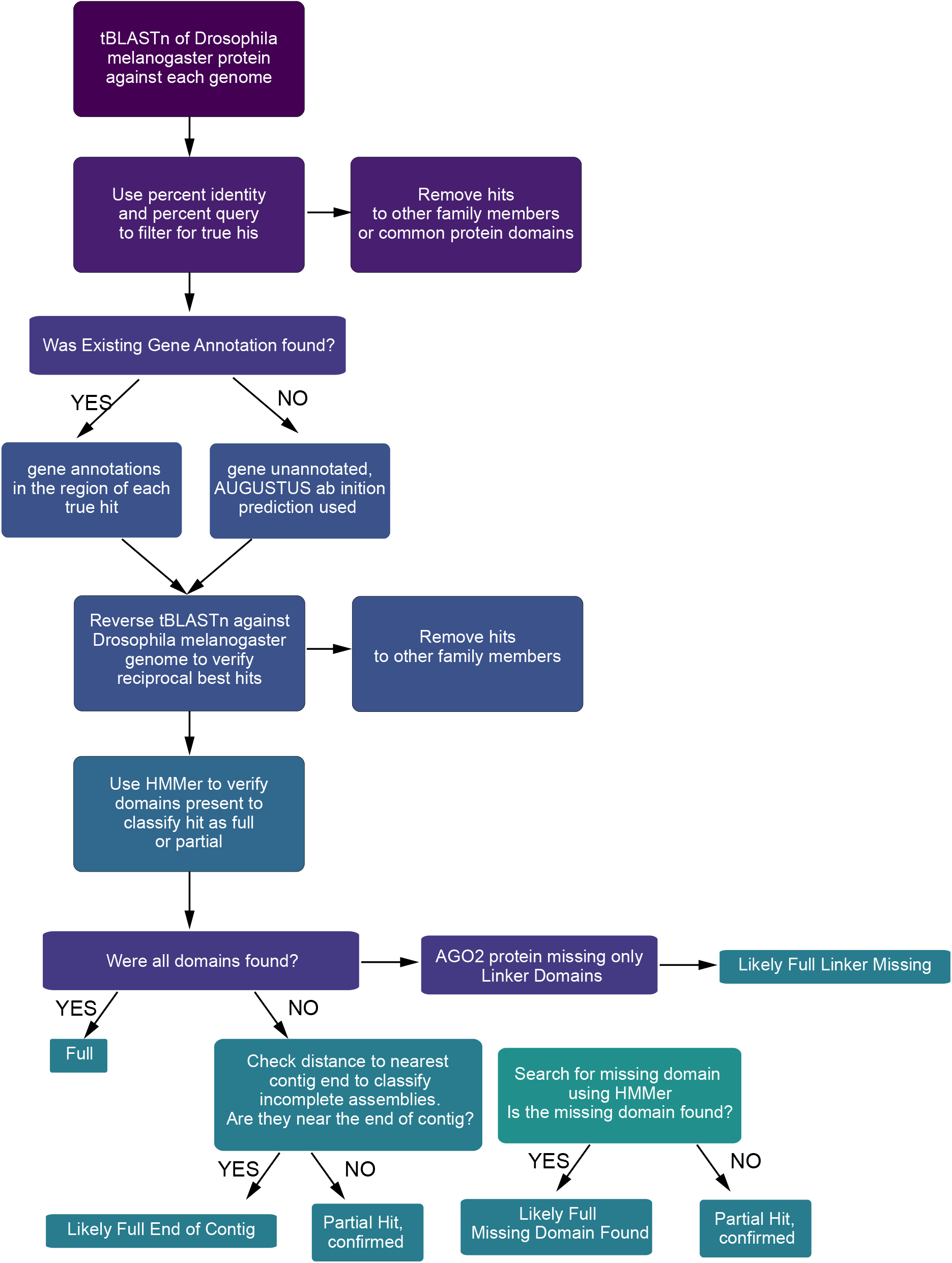
Pipeline for identifying core small RNA machinery across >300 Drosophilid genomes. Recognizing that repetitive and fast-evolving gene families pose challenges for automatic genome annotation strategies, we elected to utilize an approach that involved substantial manual curation to classify and refine gene models, and appropriately assign them as orthologs and/or paralogs of specific small RNA factors from *D. melanogaster*. We incorporated RNA-seq data where possible, but we did not require expression evidence for our annotations, recognizing that exon boundaries may not be accurate in these cases.

We assumed a certain amount of assembly artifacts to confound the accuracy of these searches. For example, genes at the end of contigs would be erroneously split into multiple partial copies; and as many of the genomes came from wild individuals, there is the possibility of the two alleles to be artifactually included in the assembly. We expected to be able to set a baseline on such artifacts via initial searches of core miRNA factors. These evolve under strong negative selection and the core fly factors share 1-to-1 orthologs in humans. Here, we searched for the nuclear RNase III enzyme Drosha and its dsRBD partner Pasha, the cytoplasmic RNase III enzyme Dicer-1 and its dsRBD partner Loquacious (Loqs), and the miRNA effector AGO1; note that the specific isoform of Loqs-PD plays a distinct role in the siRNA pathway as the dsRNA processing cofactor for Dicer-2. True to expectation, we found single, apparently full-length, orthologs of these miRNA factors across nearly every Drosophilid genome inspected (**Figure 2A-B**).

**Figure 2.**
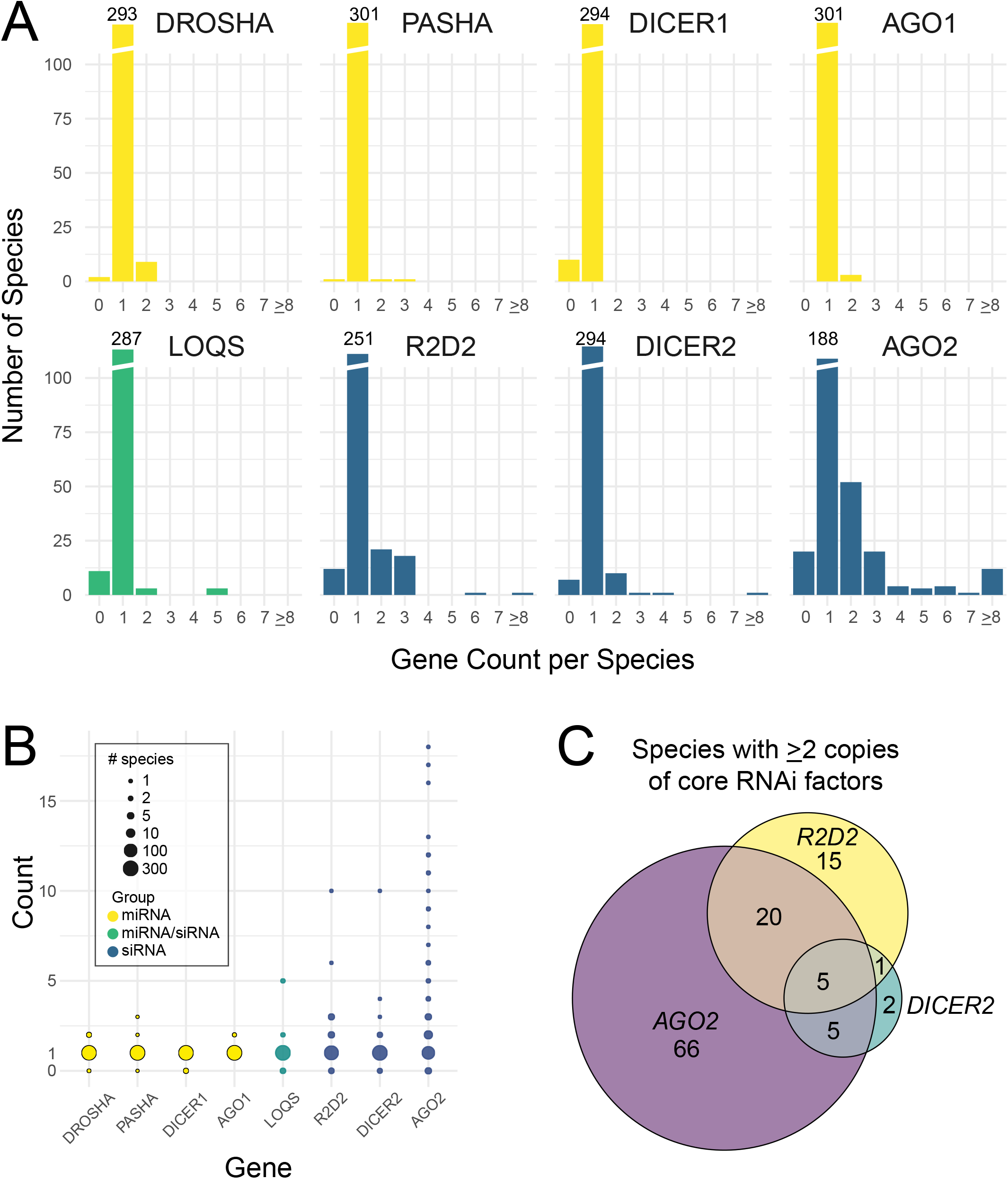
Copy number variation of miRNA and siRNA factors across fly species. (A) Top row, with only a handful of exceptions, single orthologs of all core miRNA factors were identified in nearly all genomes surveyed. We infer that many of the “non-1” instances of miRNA factor are assembly artifacts, but this sets an upper bound of 1-2% of such genome errors. Bottom row, core siRNA factors exhibit high copy number variation, with frequent duplications or amplifications in paralogs of *r2d2*, *dicer-2* and *AGO2*. (B) Dot plot emphasizing the frequency and magnitude of RNAi factor copy number changes. Overall, close to half of surveyed Drosophilid species harbor copy number variation in core siRNA factors. (C) Venn diagram shows that the majority of species bearing supernumerary copies of *r2d2* or *dicer-2* also contain one more extra copies of *AGO2*. Five species contain extra copies of all three core RNAi factors.

Closer inspection of species with “non-1” values for core miRNA factors revealed some predictable issues with these draft genomes. Even though they are based on nanopore long reads [26, 27], the contiguity of the assemblies varies across the hundreds of genomes. For most of the cases where we did not recover a full-length ortholog, we observe specific loss of N- or C-terminal domains, and find that the hits lie at the 5’ or 3’ end of a contig. Therefore, we assume that these are simply truncated or broken gene copies. If we assume that these are relatively evenly distributed across genomes, we estimate that ∼1% of cases would be truncated or missing due to genome assembly issues.

We draw attention to three potentially interesting cases of miRNA factor copy number change. First, *D. miranda* is an isolated case of an obscura clade species with extra *drosha* and *AGO1* genes. Second, a subset of related montium subgroup species (*D. lacteicornis*, *D. asahinai*, *D. rufa* and *D. tani*) all exhibit an additional copy of *drosha* (**Figure 3A**). Third, a subset of highly related quinaria group species exhibits additional copies of several miRNA factors. Puzzlingly, these affect different genes: *D. subquinaria* with an extra *drosha* copy, *D. suboccidentalis* with an extra *pasha* gene, and *D. recens* with duplications of both *drosha* and *AGO1* (**Figure 3A**). Since these species are highly related, but duplicate different miRNA factors, but other sister species lack additional miRNA factor copies, it seems that this state is in high flux.

**Figure 3.**
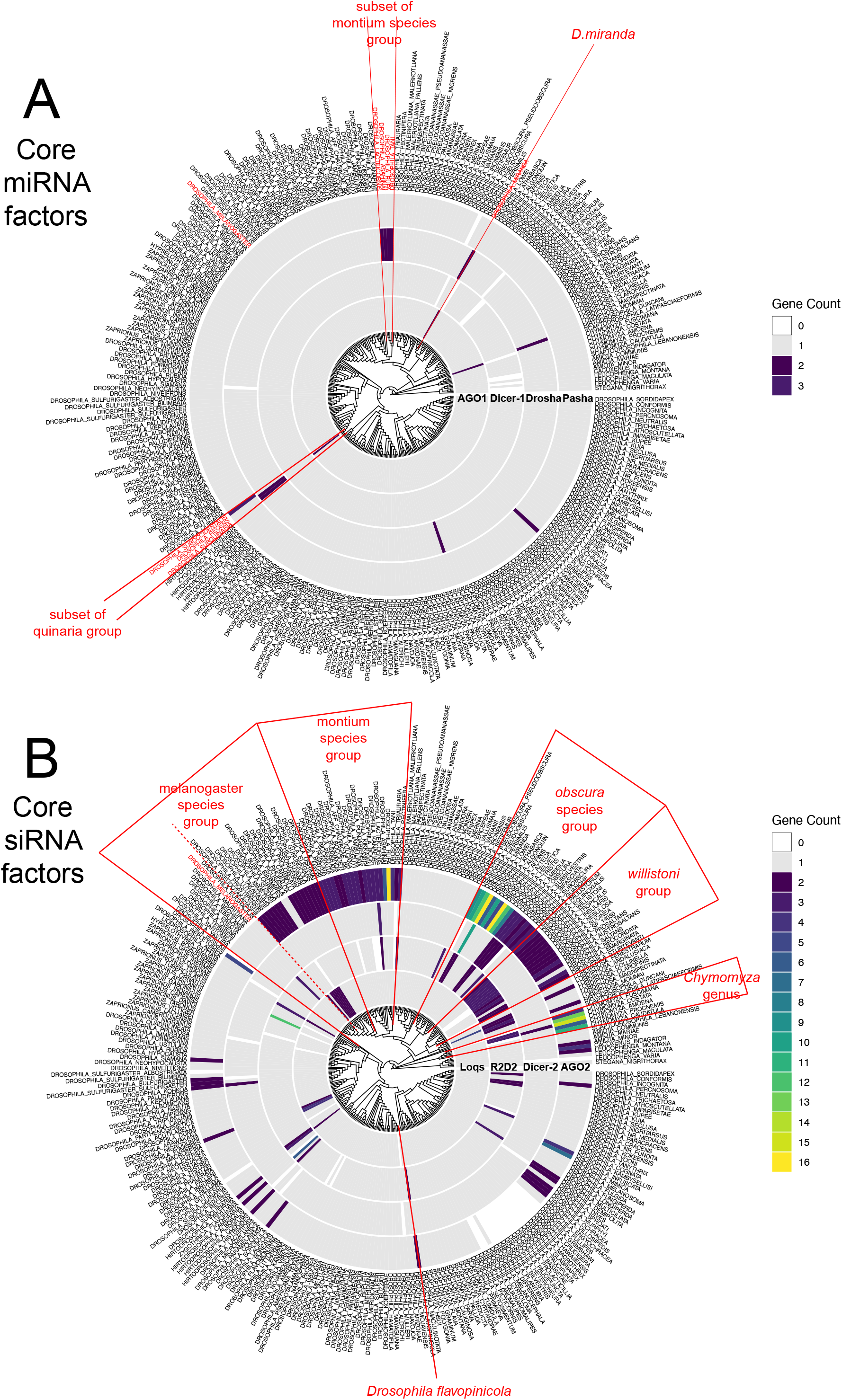
Copy number variation of core miRNA and siRNA factors by phylogeny. Circos plots illustrate the evolutionary relationships and co-occurence of copy number changes in small RNA factors across species. (A) As noted, miRNA factors are highly stable and generally exist in a single copy across the vast majority of species. The small number of species with putative changes, especially lack of a given ortholog, may potentially reflect genome assembly artifacts. A few provocative cases are shown in which related species harbor additional copies of core miRNA factors, as in a subset of montium group and quinaria group species. D. miranda is also highlighted, since it has extra copies of both *drosha* and *AGO1*. (B) siRNA factors exhibit high frequencies copy number changes. Of these *AGO2* is expanded in the most species (96/304), followed by *r2d2* and then *dicer-2*. Many clades of Drosophilid species clearly share core siRNA factor amplifications, presumably reflecting expansion in an ancestral species. This is especially striking within the willistoni clade, for which most species contain additional copies of both *AGO2* and *r2d2*. The obscura clade is notable for harboring the largest numbers of *AGO2* amplifications (visible as sectors in green shades or yellow). Subsets of obscura clade species also bear *r2d2* or *dicer-2* expansions. Finally, many sporadic, evolutionarily recent, expansions of core siRNA factors can be observed across the Drosophilid phylogeny. The example of *D. flavopinicola* is highlighted as all other species along its phylogenetic branch lack RNAi factor expansions, but it has extra copies of both *r2d2* and *AGO2*.

### Recurrent duplications and amplifications of RNAi factors across Drosophilid genomes

With the analytical framework in hand, and with knowledge of underlying issues when working with the Drosophilid genomes, we next tackled RNAi factors. Since copy number changes in *AGO2* were documented in collective studies of ∼40 fly species [23–25], we expected to be able to identify more cases when sampling nearly an order of magnitude more species. Indeed, we were not disappointed in this endeavor.

We find 96 species with extra copies of *AGO2*, ranging from 2 up to 16 copies. When arranging these *AGO2* expansions by phylogeny, we find numerous cases of ancestral amplification within various subclades that span the Drosophilid family tree. Several large groups of related species harbor extra *AGO2* genes, along with numerous isolated species or pairs of sister species that harbor an extra *AGO2* copy (**Figure 2A-B**). Beyond the established case of the obscura clade, other notable subgroups include the willistoni and montium species, within which most members exhibit extra *AGO2* copies. Interestingly, within these three major clades of ancestral expansion of *AGO2* genes, we can clearly observe sub-lineages with accelerated dynamics of *AGO2* amplification. For example, amongst the montium clade species, several members within montium subgroup (e.g. *D. tani*, *D. asahinai*, *D. rufa*) specifically amplified their *AGO2* paralogs to 5, 6, and 11 full length copies, respectively. Likewise, amongst the obscura clade species, the *pseudoobscura* sublineage exhibits the largest amplifications of AGO2 genes, including *D. pseudoobscura pseudoobscura* (>9), *D. pseudoobscura* (10), *D. persimilis* (>12), *D. miranda* (>16) and *D. lowei* (6). These observations appear to reflect recurrent, escalating arms races, that involve RNAi effectors. At the other extreme, we also find examples of very recent expansions of *AGO2* genes. For example, the related Hawaiian species *D. kambysellisi* and *D. infuscata* harbor at least 4 and 7 full-length AGO2 genes, but the closely related sister species *D. mimica* has only one (**Figure 3B**). Overall, the clear evolutionary relationships amongst numerous species that bear expanded *AGO2* genes lends strong credence to the notion that these are genuine copies, and not assembly artifacts.

To date, copy number variation of other core RNAi factors has not been reported in *Drosophila* species, with the exception of amplification of *dicer-2* (and *AGO2*) on the neo-sex chromosomes of *D. miranda* [29]. As noted, we found a few species that candidately harbor extra copies of *loqs*. However, we were surprised to find several species with extra copies of *dicer-2*, and >40 species with duplications of *r2d2* (**Figure 2A-B**). Of the latter, 20 species carried 3 or more copies of *r2d2*. Species bearing expansion of *dicer-2* or *r2d2* occur sporadically across the phylogeny, but we noticed that willistoni clade species consistently harbored extra copies of *r2d2* (12 out of 13 species). Overall, the expansion of RNAi factor gene copies is a far broader feature of Drosophilid genomes than previously recognized.

### Nested expansions of RNAi factors

We noticed an apparent hierarchy of RNAi factor amplification. The high frequency of supernumerary AGO2 makes sense with the fact that levels of Argonaute factors are limiting for small RNA function [30, 31]. Thus, this is a salient lever to press when seeking the capacity for genetically enhanced RNAi. Interestingly, the copy number variations of *dicer-2* and *r2d2* do not appear randomly, since they infrequently occur by themselves and instead typically co-occur with *AGO2* expansion (**Figure 2C**). We also take notice of the fact that amongst Dicer-2 dsRBD partners, there is a predilection for R2D2 (but not Loqs) to expand. Dicer-2 partners with the isoform Loqs-PD for siRNA biogenesis [32–34], but partners with R2D2 to load siRNAs into AGO2 [35, 36]. The fact that either Dicer-2 or R2D2 tends to be elevated in copy number may imply that efficient siRNA loading is critical when modulating endo-siRNA capacity during evolution.

The phylogenetic relationships of species bearing copy number changes in small RNA factors are easier to appreciate using circos plots. As mentioned, a couple pockets of related Drosophilid species harbor coordinated changes in core miRNA factors, which deserves future investigation (**Figure 3A**). However, the richness of phylogenetic diversity in core RNAi factors across ∼300 Drosophilids is stunning (**Figure 3B**). A notable example of RNAi factor co-occurrence exists within the willistoni group, for which 12/13 species harbor multiple *AGO2* and *r2d2* genes. Therefore, it appears that there was co-expansion of multiple RNAi factors in a willistoni ancestor, which has been propagated into most of the present-day descendant species (**Figure 3B**). A distinct set of evolutionary dynamics is illustrated by the montium group species. Nearly all of these exhibit expansion of *AGO2* genes, but at the tips of this subphylogeny, two species independently amplified *r2d2* (*D. pectinifera* and *D. kanapiae*); the latter has further expanded *dicer-2* (**Figure 3B**). Finally, yet another set of complicated trajectories is illustrated by the *obscura* clade, whose species universally harbor extra copies of *AGO2* (**Figure 3B**).

Amongst these, we can discern that AGO2 genes themselves have amplified more radically along one sublineage of obscura clade species that includes *D. pseudoobscura*, *D. persimilis* and *D. miranda*, and others, as noted above. However, we also observe different sublineages of obscura clade species that have amplified *dcr-2*, or that have amplified *r2d2*, although none had specifically amplified *loqs*. Amongst the remainder of the Drosophilid phylogeny, we observe numerous sporadic expansions of particular RNAi factors (**Figure 3B**). Given the paucity of such events for core miRNA factors (**Figure 3A**), these collectively indicate that acquisition of additional RNAi factor gene copies is an extremely recurrent event. We highlight *D. flavopinicola*, as it resides along a deep branch of the Drosophilid phylogeny for which none of the other species harbor additional RNA factors. Strikingly, *D. flavopinicola* exhibits extra copies of both *r2d2* and *AGO2*, suggesting a highly focal need to amplify RNAi pathway function in this species. Finally, we note five species exhibit the “jackpot” and confidently harbor multiple copies of all three core RNAi factors (**Figure 3B**). Three of these reside within a sublineage of obscura clade species (*D. miranda*, *D. tristis*, and *D. ambigua*), but the others (*D. kanapiae and Chamomyza amoena*) are evolutionarily unrelated events.

To further highlight these dynamic and coordinated changes of copy number variation in miRNA and siRNA factors, we plotted these data with respect to several specific Drosophilid sublineages mentioned above. These plots illustrate examples of atypical expansion of core miRNA pathway factors (**Figure 4A-B**), but more generally illustrate the diversity of evolutionary dynamics for core siRNA pathway factors (**Figure 4A, C, D**). In particular, the different nested amplifications are consistent with the notion that they secondarily support enhanced RNAi activity within pre-existing situations of AGO2 expansion. Taken together, these evolutionarily independent examples of coordinated expansions of RNAi factors conform well to the scenario of recurrent escalating genetic arms races, in which increasing stockpiles of different core RNAi factors are likely used to fight different intragenomic conflicts.

**Figure 4.**
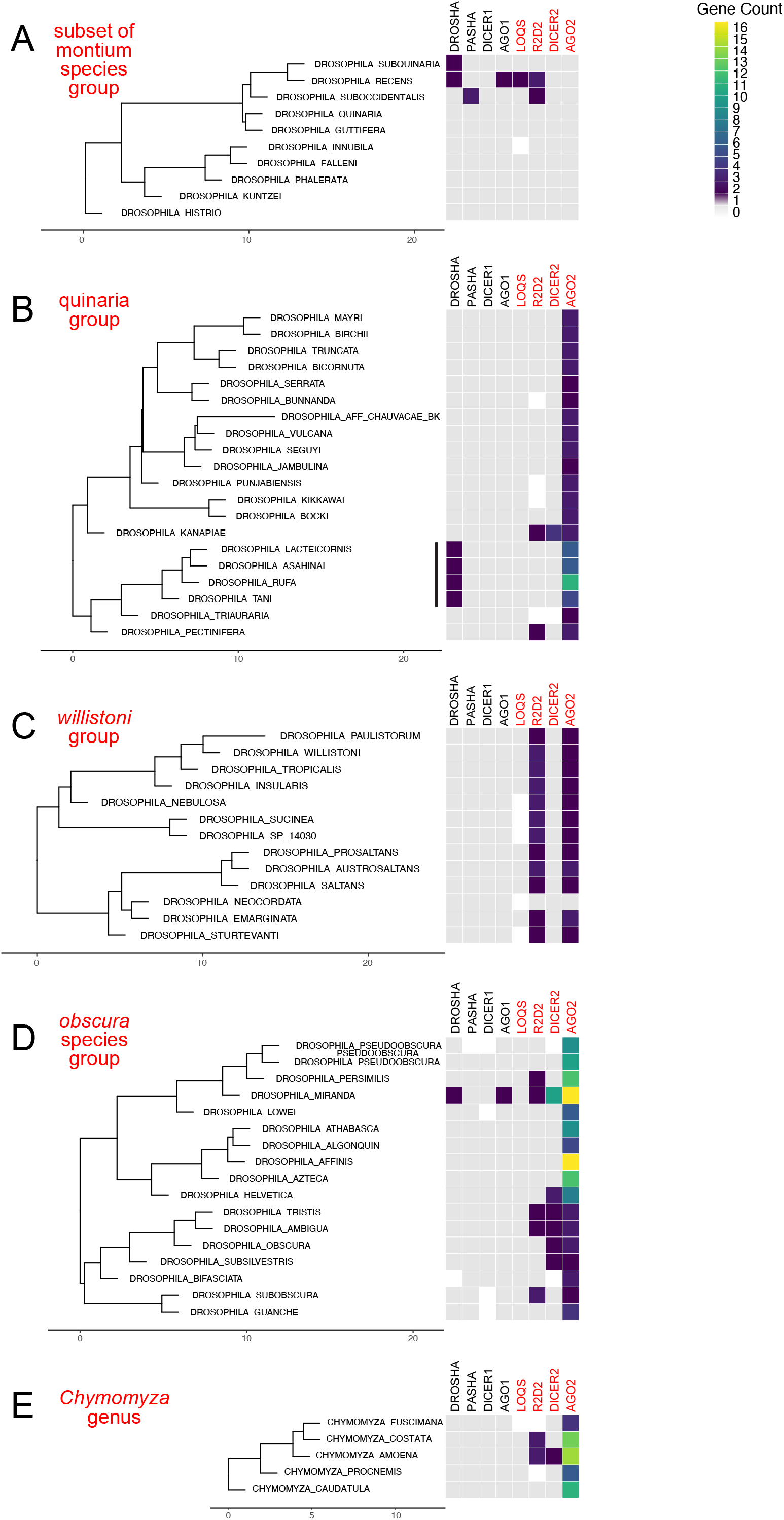
Examples of ancestral and/or nested expansions in core siRNA factors. (A) A subset of montium group species have expanded both the core miRNA factor *Drosha*, as well as the siRNA effector *AGO2*. (B) A sublineage of the quinaria clade has expanded several members of the core miRNA pathway, and occasionally some siRNA factors. Curiously, the nature of the small RNA factor changes does not clearly align with phylogenetic relationships. (C) Nearly all species across the willistoni group have co-amplified siRNA factors *r2d2* and *AGO2*. (D) All species within the obscura clade have amplified *AGO2*, with subsets of species also gaining copies of the other siRNA factors. Apparent evolutionary trajectories can be inferred from the phylogeny. For example, in the lower half of the obscura phylogeny, which all exhibit amplification of AGO2, *D. subsilvestris* and its sister species all gained *dicer-2*, while *D. tristis* and *D. ambigua* further gained *r2d2*. (E) All species analyzed within the *Chymomyza* genus have amplification of *AGO2*, and a subset of them have additional expansion of *dicer-2* and *r2d2*. Note examples of species within the obscura group and *Chymomyza* that have expanded all three core RNAi factors.

### Testis-biased expression of RNAi factors, including expanded copies

It is known that several expanded copies of Drosophilid *AGO2* are preferentially expressed in testis [24, 25]. We examined this further using a broad set of RNA-seq data from ovaries and testes [37]. Because we wished to be certain on the provenance of reads across paralogs, we utilized only uniquely mapping RNA-seq data for this analysis. These efforts confidently detected expression of the strong majority of RNAi copies that we annotated (**Figure 5A**). Of note, when comparing the expression of core miRNA factors with RNAi factors, we generally observe that miRNA factors are higher expressed in ovaries compared to testis (**Figure 5B**). We do not interpret this to imply preferential function of miRNAs in the female germline; rather, it more likely reflects the provisioning of miRNA factors into the oocyte. This provides a baseline expectation that some presumed generally utilized genes are not equally expressed between gonads, which is fully expected from the biology of maternal deposition that fuels initial embryo development.

**Figure 5.**
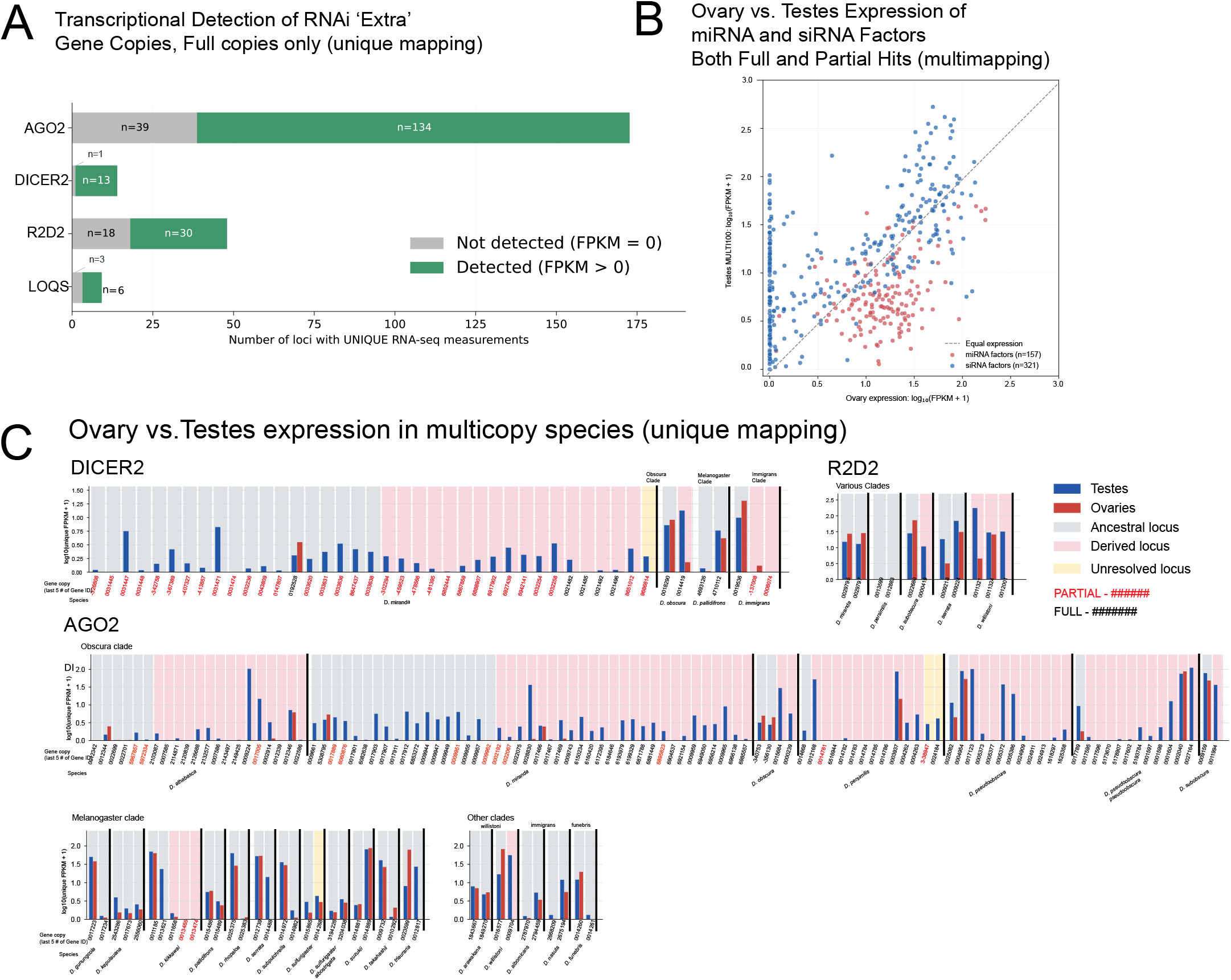
Testis-biased expression of many RNAi factors. We analyzed the expression of core small RNA factors across numerous species with paired ovary and testis RNA-seq data. (A) By conditioning on uniquely mapping reads, we could verify the expression of the strong majority of expanded RNAi factor copies annotated in this study. (B) A general trend across species is that miRNA factors are preferentially expressed in ovaries, likely for maternal provisioning, while siRNA factors are generally biased for preferential expression in testis. (C) Gonadal expression patterns of individual RNAi factors across selected species with expanded copies.

With this in mind, it seems notable that there is clearly opposite sex bias to the vast majority of siRNA factor copies we annotated across fly species, with most of them preferentially expressed in testis (**Figure 5B**). This was confirmed by analyzing these at the level of individual copies, where we found numerous derived copies of *dicer-2* and *AGO2* were preferentially expressed in testis (**Figure 5C**). These trends were particularly the case for species within the obscura clade. However, we note that testis-bias was not universally true across all species harboring RNAi copies.

Overall, this analysis extends prior observations on the preferential deployment of core RNAi factors and/or specialized paralogs in testis. While this was known for AGO2 [24, 25], to our knowledge this is the first recognition of recurrent trends for enhanced and/or expanded utilization of Dicer-2 in testis.

### Rampant emergence of *de novo* hpRNA loci in *D. pseudoobscura*

We and others characterized inverted repeat transcripts, termed hairpin RNAs (hpRNAs), as a major source of *D. melanogaster* endogenous siRNAs with trans-acting capacity [38–40]. We subsequently recognized that all *D. melanogaster* hpRNAs evolved recently [14], and therefore do not mediate conserved gene regulation. Instead the rapid dynamics of hpRNA birth and death [41] befits their genetic requirement to suppress rapidly evolving meiotic drivers [15, 16] or suspected meiotic drivers [5, 14].

Given the amplification of RNAi factors across most obscura clade species (this study and [23–25]), we wondered whether hpRNAs might exhibit notable evolutionary dynamics in this clade. We selected *D. pseudoobscura* for this effort, owing to its wealth of existing small RNA and RNA-seq datasets [42–45]. Importantly, many of the small RNA datasets are from testis, the known dominant location of siRNA accumulation in characterized fly species, but other available tissues included male body, female body, head, ovary, imaginal discs and embryos. We mapped these datasets and used previous strategies to manually curate transcripts bearing inverted repeats, whose arms generate substantial numbers (>1000 reads) of characteristic siRNA-sized (predominantly 21-22 nt) reads [5, 14] (**Figure 6A**).

**Figure 6.**
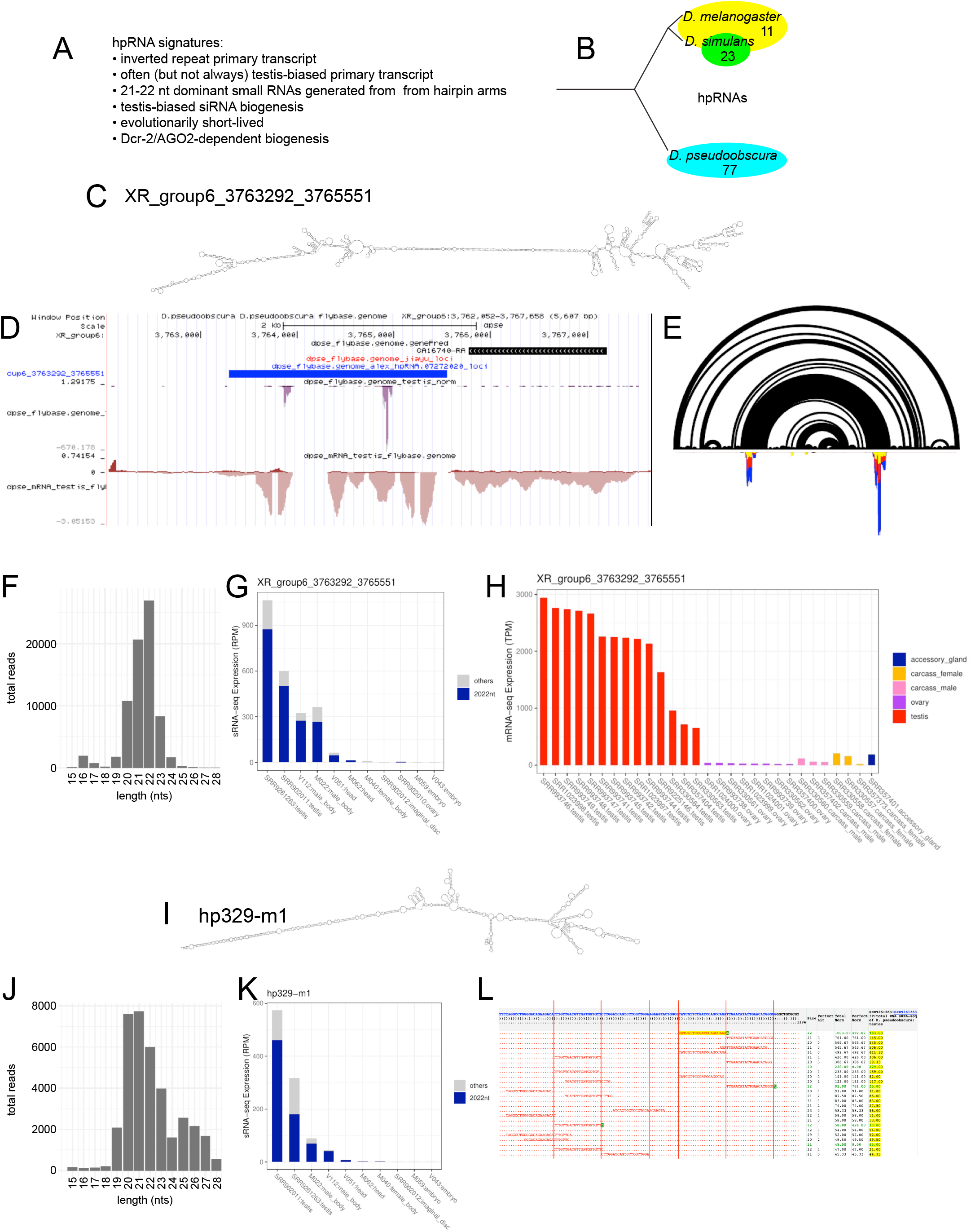
Annotation of *D. pseudoobscura* hpRNAs. (A) General features of hpRNAs that we used for annotation using *D. pseudoobscura* small RNA data from several tissues, including testis. Note that we cannot use genetic criteria to assess *D. pseudoobscura* hpRNAs, as CRISPR-mediated mutagenesis is extremely challenging in this species. (B) Numbers of confidently identified hpRNAs in three *Drosophila* species. The close sister species *D. melanogaster* and *D. simulans* share 11 hpRNAs, but the later has nearly two dozen species-specific hpRNAs. By comparison, *D. pseudoobscura* has far more hpRNAs than either of these, and has no loci in common. (C-H) Example of newly annotated *D. pseudoobscura* hpRNA, located at XR_group6_3763292_3765551. (C) Overall hairpin structure. (D) Transcriptome data reveals an unannotated, splice transcript expressed in testis, which is associated with prominent small RNA peaks. (E) Comparison with an arc plot (top) shows that the small RNAs (bottom) are specifically associated with the duplexed regions of the locus. (F) Small RNAs from this locus are predominant 21-22 nt in length. (G) Small RNAs from this hpRNA are preferentially found in testis libraries. (H) The primary hpRNA transcript is also predominantly expressed in testis. (I-L) Example of newly annotated *D. pseudoobscura* hpRNA, hp329-m1, which forms an extended hairpin (I). (J) Its small RNA profile is biased for 21-23 nt species reflecting siRNAs, but also seems to have a tail of piRNA-sized species (25-27 nt), which are testis-restricted (K). The dominant small RNAs from the hp329 hairpin are specifically siRNA-sized, and clearly reflect phased biogenesis of consecutive RNase III cleavages.

Considering that *D. melanogaster* has only 11 hpRNAs [5], these efforts yielded a bonanza of hpRNAs in *D. pseudoobscura*. As in our previous small RNA annotation efforts, we sought to separate “confident” hpRNAs with multiple clear features, from candidate loci (which did not pass one or more cutoffs). We note that the latter class may still include genuine hpRNAs, and may become confident with deeper sequencing and/or *dcr-2* mutants, which do not yet exist in this species. Nevertheless, we classified 77 confident hpRNAs, with 29 additional candidates (**Figure 6B**). Given that hpRNAs are rapidly-evolving, we were not surprised to see that all of our newly annotated loci were unique to *D. pseudoobscura*. We take the fact that this species harbors so many more hpRNAs than either *D. melanogaster* or *D. simulans* as a strong signal for its heightened usage of endogenous RNAi, going hand-in-hand with its amplification of the RNAi effector Ago2.

We illustrate the collective properties of this enormous set of hpRNAs with Dps-XR_group6_3763292_3765551 (**Figure 6C-H**) and hp329-m1 (**Figure 6I-L**) as exemplars. These hpRNAs illustrate extended inverted repeat structures within their progenitor transcripts, specific generation of siRNA-sized reads from their duplex portions, preferred accumulation of their small RNAs in testis, and tissue specificity of the primary hpRNA transcript.

Known hpRNAs differ in the specificity of siRNA production relative to their progenitor hairpins. For example, the *D. simulans* Nmy and Tmy hpRNAs [15, 16] yield heterogeneous siRNAs across the length of their duplex arms, while loci such as *D. melanogaster* hp-CG18854 and hp-CG4068 generate specific siRNAs in a phased pattern. However, while phasing is overall not very common amongst known hpRNAs, it is reasonably frequent amongst newly annotated *D. pseudoobscura* hpRNAs. For example, Dps-hp329-m1 is an example locus whose most abundant siRNAs exhibit phasing, reflecting consecutive cleavages by Dcr-2 (**Figure 6L**).

Consistent with hpRNAs we annotated in *D. melanogaster* [14] and *D. simulans* [5], nearly all *D. pseudoobscura* hpRNA-siRNAs accumulated preferentially in testis compared to other tissues (**Figure 6G, 6K**). Comparison of tissue-specific RNA-seq data showed that the majority of pri-hpRNA transcripts were also strongly testis-enriched or restricted. However, this was not always the case. Some pri-hpRNA transcripts were detected more broadly across non-testis tissues surveyed (accessory gland, ovary, male carcass and female carcass), even though the siRNAs themselves from such loci were more testis-specific. We previously observed a similar situation in *D. melanogaster* [14], and we interpret such discordance to reflect higher siRNA biogenesis activity in the male reproductive system than in other settings. Overall, these expression data affirm that, as in *D. melanogaster* and *D. simulans*, the Drosophilid hpRNA-siRNA pathway is highly testis-biased.

### Genomic clustering of *D. pseudoobscura* hpRNAs

Another feature of known hpRNAs recapitulated amongst our newly annotated *D. pseudoobscura* hpRNAs is that of genomic clustering. We previously noted that many hpRNAs exist as paralogous groups, and sometimes as tandem copies. For example, D. melanogaster hp-CG4068 harbors 20 nearly identical tandem hairpin copies. The Dps-hp551 family illustrates a newly identified multicopy hpRNA, many of which are tandemly arranged (**Figure 7A-B**). However, we also identified unexpected hpRNA arrays that contain largely unrelated hairpins, an arrangement that has not been previously recognized. For example, XL_group1e:10,073,414-10,085,239 generates a multiexonic testis transcript that includes 4 adjacent hpRNAs Dps-hp346, hp484, hp537, hp420 (**Figure 7D-E**). Curiously, sequence alignments indicated that their sequences are highly dissimilar, with share only short motifs at best, and might be incidental (**Figure 7F**). These hpRNA reads are also highly phased (**Figure 7G**). Currently, we do not know the functional or regulatory significance of clustered hpRNAs. However, it is worth noting that 30% of metazoan miRNAs are clustered, and this arrangement can accommodate specific biogenesis regimes by the Drosha/Pasha complex on an individual pri-miRNA operon transcript [1]. Similarly, It is conceivable that physical proximity of Dicer-2/Loqs-PD complexes on a single pri-hpRNA operon might have impacts on their biogenesis.

**Figure 7.**
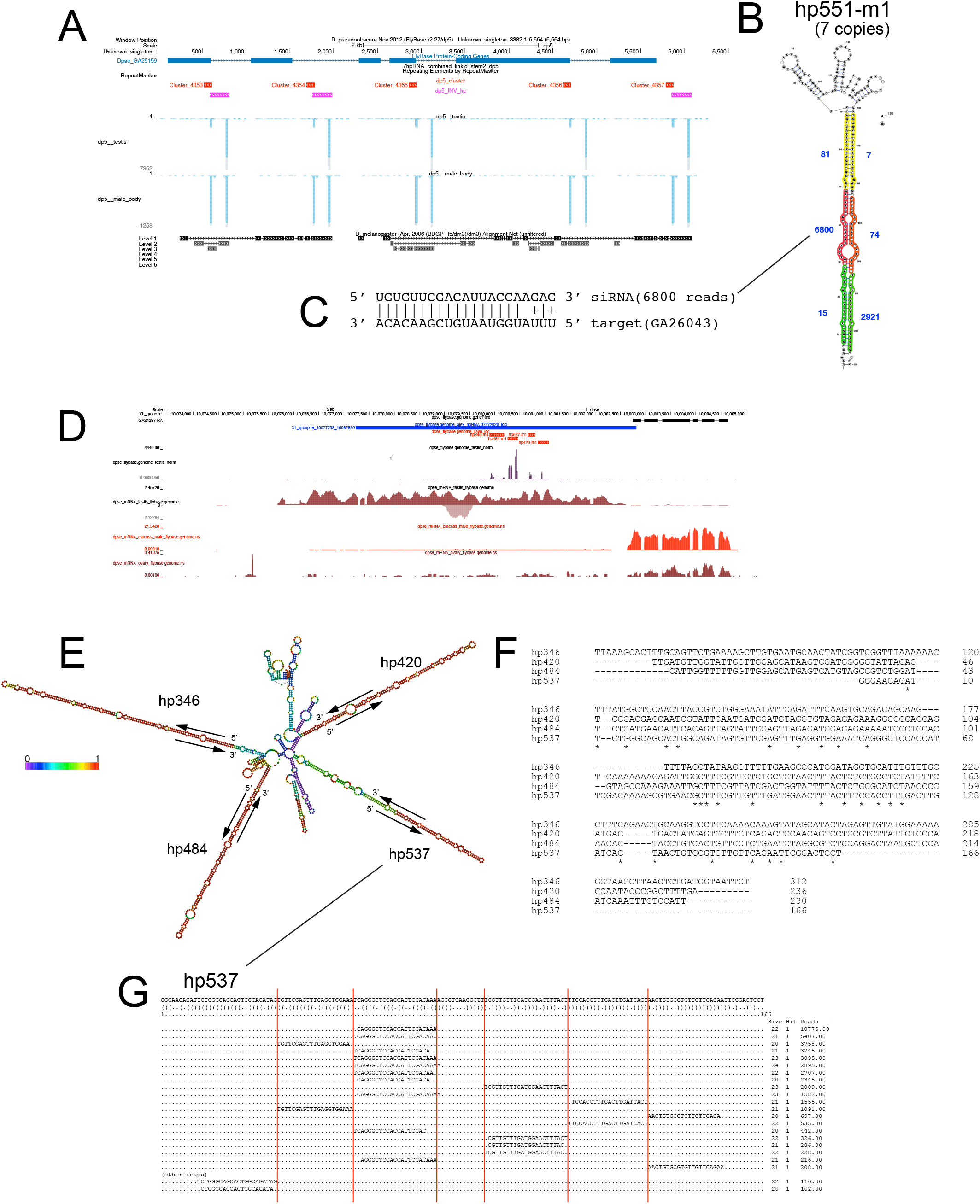
*D. pseudoobscura* hpRNA clusters. (A) Example of a hpRNA cluster composed of tandem duplicates of the hp551 family. (B) Secondary structure of one of the exact duplicate hpRNAs, emphasizing a series of predominant reads that correspond to phased RNase III cleavages, presumably by Dicer-2. (C) The most abundant hp551-siRNA has high antisense complementarity to its target gene GA26043. (D-G) Example of a hpRNA cluster composed of distinct hpRNAs. (D) This locus is defined by an unannotated transcription unit that is testis-specific; the neighboring gene is ovary-expressed. A series of adjacent inverted repeats generate specific small RNAs. (E) Predicted structure of the region bearing hp346, hp484, hp537 and hp240. (F) These hpRNAs exhibit at best only modest sequence similarity. (G) Example of dominant reads from hp537 that define phased siRNA duplexes.

We find likely target genes for a subset of its hpRNAs. For example, Dps-hp551 generates abundant siRNAs with nearly perfect antisense matching to *GA26043*, which encodes a fibronectin type III domain containing 3B (Fndc3b) protein (**Figure 6B**). Curiously, we did not identify overt targets for many *D. pseudoobscura* hpRNAs, as was typically the case in *D. melanogaster* [14] and *D. simulans* [5]. Some potential reasons might be that this species has many targets in non-assembled or polymorphic regions of the genome, harbors exuberant capacity to innovate inverted repeat transcripts independently of gene targeting, and/or might harbor relaxed rules for siRNA targeting. Relevant to the last point, it is possible that some of the numerous AGO2 paralogs in *D. pseudoobscura* may have biochemical properties that are distinct from *D. melanogaster* Ago2 [46–48], which is by far the most studied insect Slicer enzyme. As we highlighted through this work, the makeup of the RNAi pathway in *D. melanogaster* is markedly different in a large swath of other Drosophilid species, and it is not a stretch to hypothesize that at least some RNAi factors in other flies may harbor distinct activities.

## Discussion

### Intragenomic conflicts fuel arms races that recurrently involve small RNA pathways

The Red Queen hypothesis posits that species are under continual pressure to adapt and evolve, merely to maintain their status quo [49]. This is a reference to the Red Queen in Lewis Carroll’s “Through the Looking-Glass”, as this character dashes with Alice through Looking-Glass Land and comments “Now, here, you see, it takes all the running you can do, to keep in the same place”. It is well-appreciated that many instances of unusually rapid molecular evolution have to do with parasite-host relationships or situations of intragenomic conflict. Amongst these, viruses and transposons are particularly well-studied as paradigms of fast-evolving molecular conflict [50, 51]. Since deregulation of viruses and transposons are highly deleterious to organismal health and/or reproduction, numerous host strategies have been innovated to suppress these selfish elements. These rapid cycles of molecular warfare lie at the heart of Red Queen’s arms races: viruses and transposons evolve to evade suppression by host defenses, which must then innovate to keep up with these intruders.

In *Drosophila*, viruses and transposons are predominantly countered by the RNAi and piRNA pathways, respectively. Accordingly, it makes sense that many of the proteins in these small RNA pathways evolve rapidly and are subject to positive selection [52, 53], in contrast to miRNA machinery, which is highly conserved across metazoans. Still, the reasons for the rapid evolution of RNAi and piRNA factors are only partially understood. In the case of RNAi factors, this has been rationalized due to the emergence of viral suppressors of RNAi (VSRs), which can neutralize siRNA biogenesis factors or effectors [54]. This induces selective pressure for changes to the RNAi machinery. However, it is not clear that many of the positively selected sites on RNAi factors can be accounted for as binding surfaces of VSRs. Thus, there may be other reasons for their rapid evolution.

A speculative hypothesis is that some endogenous selfish meiotic drivers might also operate to antagonize RNAi machinery. Although none are recognized to do so thus far, it is clear that endogenous selfish meiotic drivers can emerge on extremely rapid timescales. For example, the *D. simulans* “Paris” sex ratio drive system is inferred to have emerged in East Africa less than 500 years ago, whereupon it spread across sub-Saharan Africa and the Indian Ocean Islands [55], but is already in decline due to intense counter-selection for resistant Y chromosomes [56]. Other molecular evolution studies indicate that numerous RNAi-suppressed genes, including putative and known meiotic drivers, emerged during recent speciation [5, 15, 16]. Given the high frequency of *de novo* hpRNA-siRNA loci in certain species that are experiencing bouts of meiotic drive, such as *D. simulans* [5] and *D. pseudoobscura* (this study), it seems plausible that some hpRNA target genes might counter the activity of the RNAi pathway itself. This proposal awaits future experimentation, for which the roadmap is provided by the discovery of these hpRNA loci.

With the piRNA pathway, it might be less apparent why piRNA proteins themselves evolve rapidly [57], since the battle itself is waged at the level of RNA sequences [52]. It is clear that many protein surfaces that connect core piRNA factors co-evolve [58], so that changes in one factor may elicit changes in others. But this does not intrinsically reveal what drives such changes. Perhaps some transposon proteins might interact with piRNA machinery. Alternatively, some TEs might turn RNA silencing against the host. For example, certain copies of the telomeric transposable element TART-A have captured a portion of *nxf2*, which may enable it to suppress this piRNA factor [59]. Again, it seems a reasonable conjecture that some meiotic drivers might also have innovated strategies to antagonize piRNA factors.

Finally, it is worth considering the rapid dynamics of these intragenomic conflicts that play out in RNA space. Of course, the extremely deep and dynamic sequencing of COVID-19 strains provided an unprecedented view into the rapid evolution of virus sequences, but this is intrinsically always occurring with viruses and phages. Infection of *D. melanogaster* by diverse, and often previously unannotated, viruses was unearthed via RNA-seq and small RNA sequencing [60, 61], including large populations of “unmappable” small RNAs that harbor characteristic 21 nt siRNA lengths [62]. Similarly, transposon content is highly idiosyncratic amongst *D. melanogaster* individuals [63], and is accompanied by highly dynamic content of piRNA clusters that encode transposon defense [7, 8].

Our current study further highlights how hpRNA-siRNA defenses rapidly turn over across *Drosophila* species. Indeed, there has been substantial innovation of hpRNA loci specifically amongst simclade species, since their recent split from *D. melanogaster* [5]. More strikingly, *D. pseudoobscura* approaches an order of magnitude more hpRNAs than *D. melanogaster*, but seems not to have any hpRNAs in common. Thus, non-coding RNAs associated with siRNA- and piRNA-mediated defenses against selfish genetic elements represent some of the most individually diverse and rapidly evolving portions of *Drosophila* genomes.

### Amplification of small RNA pathway factors as beacons for active intragenomic conflicts

It should be intuitive that cells do not have limitless flux through metabolic, signaling, or regulatory pathways. Still, until one reaches that limit, it may be difficult to gauge how far away one might be from saturating such capacity. In the early days of RNAi, it came as a surprise that sufficiently high expression of non-specific shRNAs could kill mice, due to saturation of the miRNA machinery [64]. At that time, evidence was shown that saturation of the pre-miRNA export factor Exportin-5 was responsible. Later, it was also shown that Argonaute effector proteins are ultimately limiting for the function of endogenous miRNAs and synthetic siRNAs [30, 31, 65, 66]. Indeed, there is something of a zero-sum game to silencing capacity by Argonaute proteins, in that flooding of cells of synthetic siRNAs can measurably compete with endogenous miRNAs, resulting in upregulation of their targets [67]. Accordingly, strategies that can increase the formation of Argonaute silencing complexes seem attractive for the implementation of RNAi for therapeutic purposes [68].

With respect to the historical timeline of *Drosophila* RNAi, it is notable that both its biochemical mechanisms [69, 70] and exploitation for experimental gene suppression [71, 72] occurred long before endogenous siRNAs [38–40, 73–75] and RNAi biology [14, 76] were identified. In particular, the activity of artificial inverted repeat transgenes designed to elicit RNAi [77] were recognized to be enhanced in somatic settings by overexpressing RNAi factors such as Dicer-2 [78], indicating that many tissues are suboptimal for RNAi capacity.

In retrospect, such findings from experimental manipulations can be taken as a harbinger of the fact that certain settings normally ramp up capacity for small RNA pathways by increasing the expression of core factors. A recent demonstration of this is the fact that in the *D. melanogaster* ovary, the Trailblazer transcription factor specifically elevates expression of the PIWI clade effectors Aub and AGO3, to meet the demands of the piRNA pathway to suppress *Stellate*. *Stellate* is a recently emerged sex ratio meiotic driver in testis [79, 80], and its suppression by *Su(ste)* led to the first molecular evidence for piRNAs [81]. Adaptive evolution of Trailblazer supports its capacity to induce higher levels of PIWI effectors in *D. melanogaster* compared to its closest sister species, and the need for Trailblazer can be bypassed simply by overexpressing Aub and AGO3.

A similar outcome might be achieved by increasing the copy number of Argonaute effectors. Beginning 15 years ago, it was recognized that several obscura clade species harbor additional copies of the RNAi effector *AGO2* [23–25]. We broadly extend these findings in the present study, and further identify nested amplifications of the other core RNAi factors amongst obscura clade, namely *dicer-2* and *r2d2*. Remarkably, since the earliest decades of research on fruitfly genetics, multiple obscura clade species were found to harbor sex ratio meiotic drivers. These include *D. obscura* [82], *D. pseudoobscura* [83–87], *D. persimilis* [88], *D. affinis* [84]. Equally as striking, the relevant sex ratio drive loci for these species remain largely unknown, although one of the sex ratio drivers in D. pseudoobscura is the MADF-BESS domain transcription factor Overdrive [89], and a new preprint presents evidence that X-linked amplification of the spartan protease encoded by *Gcna* may be the culprit in *D. obscura* [90].

There is abundant evidence that amongst simulans clade Drosophilid species, rapidly-evolving sex ratio meiotic drivers of the Dox family are suppressed by hpRNA-siRNA loci of the Nmy/Tmy family [5, 15–19, 91, 92]. Accordingly, we posit that there are likely direct connections amongst these burgeoning sex chromosome conflicts with expansion of the RNAi machinery. This goes hand-in-hand with our finding of a bonanza of de novo hpRNAs in *D. pseudoobscura*, whose expression and function may require increased capacity for siRNA pathway function. Alternatively, it may prove that there are subfunctionalized activities for specific paralogs of Ago2, Dicer-2 or R2D2. Of note, Drosophila testis harbors enigmatic, fast-evolving clusters of miRNAs that exhibit adaptive signatures akin to hpRNAs, which are especially abundant in *D. pseudoobscura* [93]. Distinguishing these scenarios awaits functional studies of obscura clade RNAi pathway components.

The recent finding that the *de novo* sex chromosomes of the obscura clade species *D. miranda* are associated with massive gene amplifications of multiple core RNAi and piRNA factors [29] further cements the notion that increased capacity of small RNA biogenesis and/or function is recurrently associated with settings of intragenomic conflict, especially between sex chromosomes. We propose that many species we documented to harbor extra copies of RNAi and piRNA factors, many detected within new subclades of the Drosophilid phylogeny, may currently be engaged in molecular warfare involving small RNAs. According to known examples, it seems likely that many of these are involved in the suppression of selfish endogenous genes, perhaps sex ratio meiotic drivers, by siRNAs and/or piRNAs. However, we should also keep an open mind that some of these adapted copies of small RNA factors might not serve a wildtype function. It will be intriguing to characterize this tranche of novel small RNA gene copies, to see if any were co-opted for nefarious reasons by selfish genetic systems to evade silencing, as dominant negative or neomorphic small RNA machinery.

In any case, it seems highly worthwhile to peek under the hood of many of these species harboring expansions of small RNA factors, for evidence of associated intragenomic conflicts. Time and research will reveal whether the amplification of small RNA pathway machinery is generally a harbinger of genomic duress.

## Materials and Methods

### Identification of small RNA pathway genes across Drosophilid species

Small RNA pathway genes were analyzed across 304 Drosophilid species through similarity searches of predicted protein sequences. We performed tBLASTn using core miRNA and siRNA pathway genes from *Drosophila melanogaster* as queries against available Drosophila nanopore assemblies [26, 27]. An E-value threshold of 0.01 was applied to ensure search sensitivity, and a custom output format was used to capture alignment details. For each tBLASTn hit, comprehensive percent coverage and percent identity values were calculated. Plots of percent coverage vs. percent identity were made per gene, and the clustering was used to set threshold values for each gene.

If a tBLASTn hit overlapped with an annotated gene in the Obbard annotation files [28], the annotation was used as the gene sequence. In cases where there was no annotation overlapping the hit, AUGUSTUS was used to predict the gene sequence in that region. All sequences were then reciprocal BLASTed against the *Drosophila melanogaster* genome. Only those sequences which returned the gene of interest originally queried as the top hit were kept.

### Defining full and partial small RNA pathway proteins

We then ran HMMer on the gene sequences to identify protein domains in each. Hits were classified as full if they contained all protein domains, and partial if any domains were not present. Partial protein hits that were missing exterior domains were then checked to see whether the gene was within an intron length away from the nearest end of the contig. We defined the intron length threshold as Q3 + 3×IQR, where Q3 is the third quartile and IQR is the interquartile range of the observed intron length distribution for a gene across all Drosophilid species. Genes that fall less than threshold from the end of a contig were classified as “likely full.”

### Complementation of annotations with tBLASTn

A comparison was made of the tBLASTn hit and the Obbard annotation [28]. For genes with region(s) found via BLAST not covered by the annotation, BLAST region(s) were searched for protein domains via HMMer and added to the protein sequence if it resulted in full length protein.

### Analyses of RNA sequencing data

For the obscura clade, RNA-seq data was used (where available) to confirm gene model accuracy, verify the expression of the genes and determine sex-biased expression. All RNA data was downloaded from public repositories and aligned with HISAT2 [94] using default parameters. Datasets analyzed were (testis, ovary ID): Drosophila_athabasca (SRR9967656, SRR9967645); Drosophila_pseudoobscura (SRR9225146, SRR1024000); Drosophila_persimilis (SRR7243291, SRR23592635); Drosophila_miranda (DRR055195, DRR055184); Drosophila_obscura (DRR055228, DRR055218); Drosophila_subobscura (SRR26249429, SRR31237433).

### Annotation of hairpin RNAs in *D. pseudoobscura*

We utilized published *D. pseudoobscura* small RNA datasets from the Tang lab [43]: SRR902011 (testis), SRR9261263 (testis), SRR902012 (imaginal discs), SRR902010 (ovary), and the Lai lab: [44, 45]: M022 (male body), M040 (female body), M059 (embryo), M062 (head), V051 (head), V112 (male body). We used Cutadapt to remove linkers [95] and mapped >15nt reads using Bowtie [96]. We converted BAM alignments from bowtie mapping to bigwig for visualization on genome browsers, using bam2wig.pyscript from the RSeQC package [97]. We used previous strategies to call small RNA clusters and associate 20-22 nt-biased clusters with predicted inverted repeats [5, 14]. All of the clusters were manually browsed to confirm specificity of inferred siRNA production from predicted hairpin arms, and compared to RNA-seq data to assign likely biogenesis via a single stranded primary hpRNA transcript, and to distinguish them from possible cis-NAT-siRNA and TE-siRNA origins [62]. We assigned confident hpRNA loci that generated >1000 reads. In general, confident hpRNA loci generated a majority of total reads across the entire locus that were 20-22 nt size range, although we relaxed this constraint for two loci that appeared to have overlapping piRNA content (Unknown_group_249_2783_4131 and hp337-m1).

## Acknowledgements

We thank Jeffrey Vedanayagam, Jaeah Kim and Chung-Ming Lai for helping to inspect hpRNA candidates. We thank Kevin Wei and Yasir Ahmed-Brahmiah for discussion on selection *Drosophila* species. We thank Pankaj Dhakad and Darren Obbard for discussion on annotation of Drosophilid genomes and access to their gene models. LAW lab was supported by Irma T. Hirschl and Monique Weill-Caulier Research Award and Robertson Foundation. ECL lab was supported by the National Institutes of Health (R01-GM083300, R01-HD108914, P30-CA008748). The funders had no role in study design, data collection and analysis, decision to publish, or preparation of the manuscript.

